# RIFRAF: a frame-resolving consensus algorithm

**DOI:** 10.1101/227520

**Authors:** Kemal Eren, Ben Murrell

## Abstract

**Motivation:** Protein coding genes can be studied using long-read next generation sequencing. However, high rates of indel sequencing errors are problematic, corrupting the reading frame. Even the consensus of multiple independent sequence reads retains indel errors. To solve this problem, we introduce `RIFRAF`, a sequence consensus algorithm that takes a set of error-prone reads and a reference sequence and infers an accurate in-frame consensus. `RIFRAF` uses a novel structure, analogous to a two-layer hidden Markov model: the consensus is optimized to maximize alignment scores with both the set of noisy reads and with a reference. The template-to-reads component of the model encodes the preponderance of indels, and is sensitive to the per-base quality scores, giving greater weight to more accurate bases. The reference-to-template component of the model penalizes frame-destroying indels. A local search algorithm proceeds in stages to find the best consensus sequence for both objectives.

**Results:** Using Pacific Biosciences SMRT sequences of NL4-3 env, we compare our approach to other consensus and frame correction methods. `RIFRAF` consistently finds a consensus sequence that is more accurate and in-frame, especially with small numbers of reads. It was able to perfectly reconstruct over 80% of consensus sequences from as few as three reads, whereas the best alternative required twice as many. `RIFRAF` is able to achieve these results and keep the consensus in-frame even with a distantly related reference sequence. Moreover, unlike other frame correction methods, `RIFRAF` can detect and keep true indels while removing erroneous ones.

**Availability:** `RIFRAF` is implemented in Julia, and source code is publicly available at https://github.com/MurrellGroup/Rifraf.jl

## I. Introduction

The problem of finding the consensus of a set of sequences is fundamental to bioinformatics, especially in the age of high-throughput sequencing. This paper addresses the task of re-constructing an unknown true sequence from a set of error-prone reads. Many algorithms that solve this task focus on *de-novo* or reference-guided assembly of short reads [17, 18, 15]. However, with the advent of third-generation single-molecule sequencing technologies, such as Pacific Biosciences′ SMRT sequencing protocol [6], it is now possible to perform full-length sequencing of entire genes or small genomes. Here we will focus on finding the consensus of a set of *amplicon* sequences - where the sequences have the same start and end points. An example application would be targeted sequencing of an entire gene from a viral population (eg. [11]). We focus just on the consensus reconstruction problem, assuming that reads have first been grouped by genetic identity, either using primer ID barcodes [7, 22], or some form of clustering.

Consensus sequences found via multiple sequence alignment may be inaccurate when there are few reads available, or when the reads contain many errors. SMRT sequencing in particular is known to contain mostly indel errors, especially in homopolymer runs. For example, in [11], we discovered that 80% of the sequencing errors were indels. If these indels carry over into the consensus sequence, they cause frameshift errors which corrupt the reading frame, and render the amino acid sequence uninterpretable. If a reference sequence with a trusted reading frame is available, it can be exploited to inform the consensus.

Current approaches that attempt to reconstruct in-frame consensus sequences consider these problems separately. There are approaches to infer the consensus of multiple reads, and there are approaches to correct the reading frame of an already-inferred consensus sequence. Here, we solve these problems jointly, simultaneously considering evidence from the reads and the reference sequence.

One common approach to inferring consensus sequences is from multiple sequence alignments (MSAs), from which the consensus is calculated by taking the most common base in each column. A myriad of multiple alignment algorithms are available [19], any of which may be used for this task. This paper uses `MAFFT` [9, 8] as an example of this strategy when comparing alternatives. A multitude of tools, such as the `cons` command in `EMBOSS` [20], are available for computing the consensus of these alignments. Another approach is to use a partial order alignment [13] representation of the set of sequences, and find the consensus sequence using dynamic programming to extract the heaviest bundles [12]. This paper uses `poaV`^1^ for comparison. Other implementations of this approach include `pbdagcon`^2^, which was released by Pacific Biosciences specifically for raw SMRT sequence reads, and `nanopolish` [14], which wraps `poaV2` for Oxford Nanopore reads. Finally, specialized consensus methods are available for specific sequencing technologies; these methods model the specific behavior of their target protocol, such as read length and error model. In this domain, Pacific Biosciences developed the `Quiver` [4] and `Arrow` algorithms^3^ for building circular consensus sequences from raw ZMW reads.

Existing approaches for reading frame correction (such as `FrameBot`, which we use here as a comparator) exploit frame-aware codon alignment to a protein reference, followed by inserting or deleting bases in the target sequence [24]. Related algorithms include `FALP` and `LAST` [21], `Frame-Pro` [5], `HMMFrame` [25], and others. Another approach is hybrid sequencing, which supplements long singlemolecule reads with short reads [19]. Methods such as `HGAP` [4] use hybrid sequence data to find and remove indels.

This paper introduces a new method for inferring consensus sequences of such reads: the Reference-Informed Frame-Resolving multiple-Alignment Free consensus algorithm (`RIFRAF`). `RIFRAF` considers evidence from both the reads and the reference simultaneously, allowing reads to inform the frame correction process, and is sensitive to the read quality scores to ensure that high-quality bases are more informative. These features allow `RIFRAF` to make highly accurate predictions, even for a small number of error-prone reads. Unlike other frame-correction methods, `RIFRAF` can detect true frameshift-causing indels and keep them while removing spurious indels.

## II. Methods

`RIFRAF` addresses the following sequence consensus problem. Let *t* be an unknown template sequence, which is sequenced *N* times to generate a set of *N* pairs of reads and quality scores 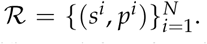 Each read *s^i^* is a noisy observation of *t*, and each *p^i^* is a vector of error probabilities, one for each base in *s^i^*. The *i*th character in read *s* is denoted *s_i_*, and the substring from the *i*th to the *j*th character is denoted *s_i_…_j_*. *p_i_* is the probability that *si* is an error; an error at a base is either a substitution, an insertion, or a deletion has occurred next to it. The task is to infer a consensus sequence *c* that matches the unknown *t*. Additionally, we also consider a reference sequence *r* and prefer that *c* not contain insertions or deletions that change its reading frame relative to *r*. This is especially useful when the template that generated the reads in 𝓡 had an intact reading frame, but the reads themselves have a high indel rate.

The structure of the full `RIFRAF` model is shown in Figure 1. It infers the unknown template by optimizing two objectives: the quality-aware alignment to the reads, and a frame-aware alignment to the reference. The optimization procedure starts with an initial consensus sequence and proceeds in an iterative greedy manner, mutating the consensus sequence at every step to improve those objectives. `RIFRAF` uses a number of techniques to speed up convergence: filtering mutations, accepting multiple mutations, forward and backward alignments, banding, batching, increasing indel penalties, and multi-stage optimization.

**Figure 1:**
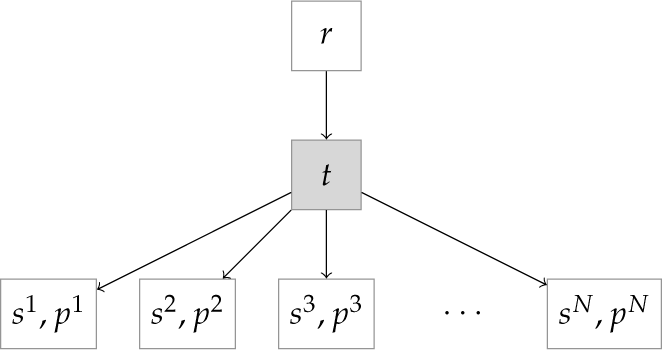
*Structure of the full model. The unknown template t (grey) has the same reading frame as known reference *r*. The sequencing process generates error-prone reads s*^1^…*s^N^ with quality scores p*^1^… *p^N^*.

`RIFRAF` is implemented in Julia [1], a high-level scientific computing language.

### i. Objective 1: pairwise alignment to reads

In order to find the optimal consensus, it is necessary to assign a score to candidates. `RIFRAF` scores consensus sequences by a global pairwise alignment [23, 16] of each read *s* with the current values of *c*. Let **A** be the |*s*| +1 × |*c*| + 1 dynamic programming matrix for aligning *c* and *s*. Each *a_i,j_* is the score of aligning prefix *s*_1…*i*_ to prefix *c*_1…*j*_. *a*_0,0_ is initialized to 0, and the last cell *a*_|*s*|+1,|*c*|+1_ contains the score for the full alignment. The score function for *c* and *s* is defined as the full alignment score: *S*(*c*|*s*) = *a*_|*s*|+1,|*c*|+1._ The overall score of consensus sequence *c* is the sum over the alignment scores for all reads: *S*(*c*|𝓡) = ∑_(*s,p*)∈𝓡_ *S*(*c*|*s*).

The sequencing process has an error rate *ρ*, which by assumption can can be partitioned into *ρ* = *ρ_mismatch_* + *ρ_insertion_* + *ρ_deletion_*. These parameters account for the different error profiles of different sequencing technologies. For instance, in SMRT sequencing, indels are more likely than substitutions. The base move scores for the alignment are derived from these error probabilities.

Typical pairwise alignment uses fixed scores for moves. However, `RIFRAF` also incorporates sequence qualities into the move scores to generate more accurate alignments. The scores for match, insertion, and deletion moves depend on the error probabilities *p* in the following way. Let *q* = log_10_ *p* (base 10 is used instead of the usual natural logarithm for compatibility with quality scores such as Phred scores). Let *q_mismatch_* = log_10*ρmismatch*_, and Similarly for the others. Then move scores are calculated as follows

- A diagonal move from *a*_*i*–1,*j*–1_ to *a_i,j_* has score log_10_ (1 – *p_i_*) if *s_i_* = *c_j_* (ie. a match), else *q_mismatch_* + *q_i_* (ie. a mismatch).
- A vertical move (insertion relative to *c*) from *a*_*i*–1,*j*_ to *a_i,j_* has score *q_insertion_* + *q_i_*.
- A horizontal move (deletion relative to *c*) from *a*_*i,j*–1_ to *a_i,j_* has score *q_deletion_* + *max*(*q_i_* + *q*_*i*+1_). If *i* = 0, the score is just *q_deletion_* + *q*_1_; similarly, *i* = |*s*|, the score is just *q_deletion_* + *q*_|*s*|_.

Intuitively, the penalties for mismatches, insertions, and deletions are more severe when the consensus does not match higher quality regions of the reads. PHRED values are capped at 30 because rarer sources of error that are not sequencing errors (eg. PCR errors) may have very confident PHRED scores, and we do not wish these to be overly informative. This cap can be adjusted if these sources of error can be ruled out (for example if PCR was not used to generate the amplicon library).

The best consensus *c*\* under Objective 1 (pairwise alignment to reads) is the one that maximizes *S*(*c*|𝓡).

### ii. Objective 2: Frame-aware alignment to reference

To perform frame correction, `RIFRAF` requires a reference nucleotide sequence *r*, which is known to be in-frame. It models the reference sequence *r* as having diverged from the template *t*, where the differences between *r* and *t* represent evolutionary events, not sequencing error as in Objective 1. The score for the consensus-reference alignment is modified to reflect this difference. First, two new moves are allowed during alignment: codon insertion and codon deletion, each with their own penalty, as shown in Figure 2. Second, a new parameter *t_indel_* is used as a multiplier for the non-codon insertion and deletion penalties. Together, these two modifications bias the alignment to prefer only codon indels, keeping the consensus in-frame. Because it uses nucleotide alignments, this method works may be expected to work better with more closely related reference sequences, where nucleotide similarity is preserved.

**Figure 2:**
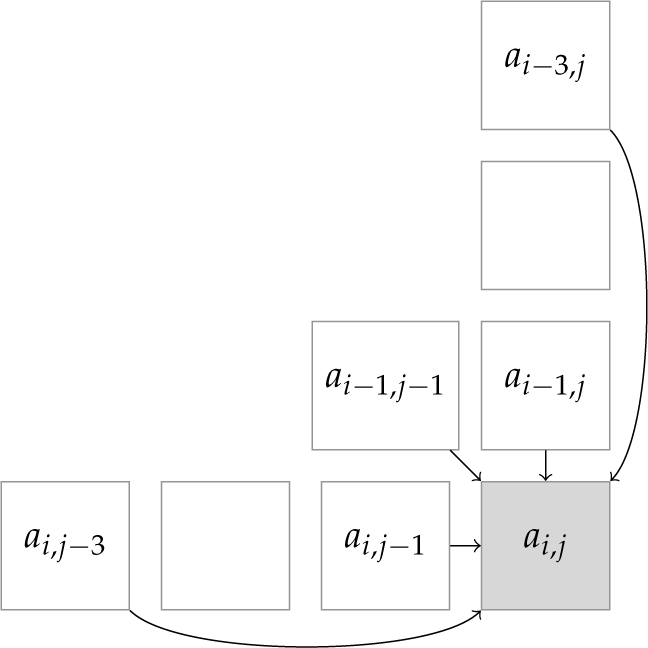
*Codon moves in the reference alignment dynamic programming matrix. The goal is to favor a consensus that preserves the reading frame. Thus, in addition to the usual single match, insertion, and deletion moves, codon insertions and deletions are also allowed, with a lower penalty than single-base indels*.

We first let `RIFRAF` converge to a draft template *c* without the reference sequence. This draft template is used to approximate the divergence between the true template and the reference, taking the edit distance normalized by the max length *d*(*r,c*)/*max*(|*r*|, |*c*|) to obtain a per-base probability of template/reference disagreement (which is used in the same manner as the per-base quality scores *p* in Objective 1). Reference (mis)match, indel, and codon error rates are provided as parameters, and the scores for each move are computed from error rate *ρ* as log_10_(*ρ*), as before.

The insertion and deletion scores are multiplied by a penalty *t_indel_*, which controls the influence of single insertions and deletions in the reference alignment. If *t_indel_* is small, frame-destroying indels may appear in the consensus, but if it is large, the consensus will be forced into the reference reading frame, even if the unobserved template really did contain indels. As we show in Section III, this penalty can be tuned to discard spurious indels while keeping true ones.

`RIFRAF` combines both objectives into a single score, allowing the reads to inform the frame correction. The score of the consensus to reference alignment is denoted *S_r_*(*c*|*r*), and the full score function is:

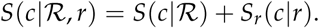

### iii. Optimization procedure

An exhaustive search for the optimal consensus *c*\* would be intractable, so `RIFRAF` uses a variant of the following greedy search algorithm, with some optimizations to speed up convergence:

1. Start with a guess *c*^0^. `RIFRAF` chooses the read with the lowest expected number of errors.
2. For the most recent guess *c^i^*, examine a set of candidate single mutations, such as insertions, deletions, and substitutions. Note that these candidates vary at each optimization stage. Keep all that improve the score *S*(*c^i^*|𝓡, *r*). Call the set of candidate mutations 𝓒.
3. If 𝓒 is empty, accept *c^i^* and terminate. Otherwise, choose some subset of 𝓒, apply them to *c^i^* to obtain *c*^*i*+1^, and iterate.

`RIFRAF` works in two stages, first optimizing just *S*(*c*|𝓡), and then optimizing the full *S*(*c*|𝓡, *r*).

#### 111.1 Filtering mutations

When comparing the template to the reads, we need not consider all possible modifications to the current consensus. For example, if any candidate mutation to *c* does not appear in any pairwise alignment of c with a read, that mutation need not be scored. Since it has no support among any observed sequence, it is likely to hurt the alignment score. Similarly, during the frame correction stage, the model only proposes insertions or deletions that appear in the pairwise alignment to reference.

#### 111.2 Multiple mutations

Instead of accepting only the best mutation in 𝓒, `RIFRAF` accepts all the mutations that are separated by a certain number of positions: *n_separate_* (the default value is 15, i.e. five codons). The candidates are accepted in order from best to worst score. This policy allows `RIFRAF` to converge in many fewer iterations than if it only accepted one mutation per iteration. *n_separate_* ensures that the changes to the consensus are relatively independent of each other, and that the score of one is unlikely to be affected by the acceptance of another. After accepting mutations in 𝓒, `RIFRAF` also compares the new score to the score that would be obtained from accepting only the single best mutation in 𝓒, and optionally accepts that single mutation instead if it results in a better score.

#### iii.3 Forward and backward alignments

Recomputing the full alignment matrix for each candidate mutation to *c* would be prohibitively expensive. For a sequence *c* from alphabet {*A, C, G, T*}, there are 4(|*c*| + 1) insertions, 3|*c*| substitutions, and |*c*| deletions to consider. Computing the alignment matrix **A** for each candidate requires *O*(*cs*) operations, so each iteration of the proposed algorithm would require *O*(*Nc*^2^*s*) operations (we omit | · | in *O*(·) for clarity). Instead, `RIFRAF` uses forward and backward alignments to compute the new score for any single change to *c* by only recomputing a single column of **A** [4].

To achieve this, in addition to the prefix alignment matrix **A**, where *a_i,j_* is the score for aligning prefix *s*_1…*i*_ to prefix *c*_1…*j*_, `RIFRAF` also computes the suffix alignment matrix **B**, where *b_i,j_* is the score for aligning suffix *s*_*i*+1…|*s*|_ to suffix *c*_*j*+1…|*c*|_. Note that *a*_|*s*|,|*c*|_ = *b*_0,0_ is the score for the full alignment. For any *j*, that alignment score can also be computed from columns **A**.,_*j*_ and **B**_.,*j*_:

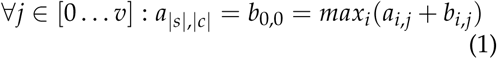

Modifying *c*j** leaves unchanged columns 0…*j* – 1 of **A**, and also leaves unchanged columns *j*… *ν* of **B**. Therefore, for all three types of mutations, computing the new score requires that at most only a single new column of **A** must be recalculated.

1. substitution at *c_j_*: compute **A**_.,*j*_; new score is *max_i_* (*a_i,j_* + *b_i,j_*).
2. insertion after *c_j_*: compute **A**_.,*j*+1_; new score is *max_i_*(*a*_*i,j*+1_ + *b_i,j_*).
3. deletion of *c_j_*: no new column necessary; new score is *max_i_*(*a*_*i,j*–1_ + *b_i,j_*).

Using the forward and backward alignments, all possible mutations to the consensus can be scored in *O*(*Ncs*) operations.

During the alignment of the template and reference, additional columns must be recomputed to account for codon insertion and deletion moves.

#### iii.4 Banding

Despite the improvements from using forward and backward alignments, each iteration is still approximately quadratic in the length of the consensus, assuming |*c*| ≈ |*s*|. Alignment banding [2, 3] further reduces the number of operations per iteration. For a given bandwidth parameter *b*, the maximum usable column size in **A** and **B** is 2*b* + ||*s*| – |*c*|| ≪ |*s*|, so evaluating a possible mutation requires many fewer operations than recomputing the full column. Alignment moves are only allowed to originate inside the band, so alignment paths must stay within the band boundaries (see Figure 3). With banding, the time complexity per iteration becomes 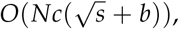 since ||*s*| – |*c*|| grows like 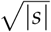 under reasonable assumptions.

**Figure 3:**
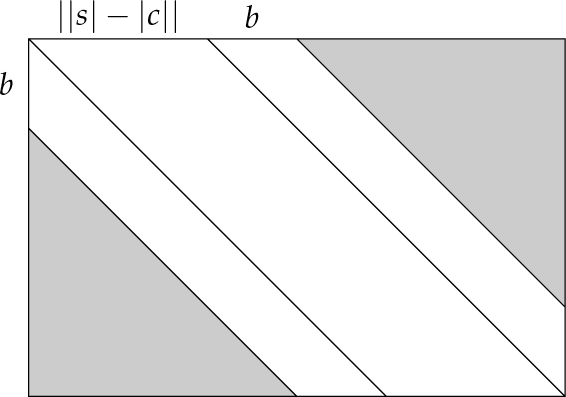
*Banded alignment. Alignments must stay within the banded region of the dynamic programming matrix*.

`RIFRAF` dynamically increases the bandwidth if the number of differences in the banded alignment is sufficiently larger than the expected number of differences implied by the read's quality scores, under the assumption that the difference between the template candidate *c* and the true template is much smaller than the number of sequencing errors in *s*. Let *r* be the observed number of differences between *s* and *c*, and *e* be the expected number of errors computed from the quality scores *p*. If the value of *r* is in the upper tail of a Poisson distribution with mean parameter *e*, then the bandwidth is doubled and the alignments are re-computed. *α* controls the size of this upper tail probability, with a default value of 0.1.

#### iii.5 Batching

`RIFRAF` uses a variety of batching strategies to speed up convergence. If the number of reads is greater than a threshold *k* (default 5), the best *k* reads by error rate are fixed as the initial batch, and `RIFRAF` runs to convergence. This ensures that `RIFRAF` first converges without considering the many spurious mutations presumably present in less accurate reads. The resulting initial guess is further refined at the refinement stage, this time with a different random batch of size *n_batch_* (default 20) for each iteration. Sequences are chosen for inclusion in the batch by sampling from a multinomial distribution of their error rates, parameterized by parameter *ρ* between 0 and 1. When *ρ* = 1, all the weight is evenly distributed among the top *n_batch_* sequences. Interpolating from *ρ* = 1.0 to *ρ* = 0.5, the probabilities become proportional to the read error rates. Interpolating from *ρ* = 0.5 to *ρ* = 0, the probabilities become uniform. By default, *ρ* = 0.9^*i*^, where *i* is the number of iterations since random batching activated. Like the fixed batch, this strategy speeds up convergence by initially avoiding inaccurate reads, then gradually letting them contribute to resolve uncertain bases if necessary.

Ideally, *n_batch_* is small enough to make each iteration fast, but large enough that RIFRAF converges stably. RIFRAF tries to detect if *n_batch_* is too small by monitoring the change in score after each iteration. If the new score is worse than the old score by more than a certain percent (10% by default), *n_batch_* is increased to 2*n^batch^*, then 3*n^batch^*, etc.

Combining all of the previous optimizations, a single iteration's time complexity is reduced from 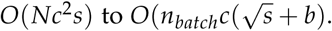

### iv. Increasing indel penalties

Whenever the algorithm converges to a consensus *c^i^*, if single indel moves were used in computing *S_r_* (·), the single insertion and deletion scores are multiplied by a parameter *t_indel_*, increasingly encouraging the alignment with the reference to use only codon indels, thereby keeping *c* in-frame. This process repeats up to *m* times, so the maximum multiplier is (*t_indel_*)^*m*^. If the penalty is large enough, the consensus will always be forced into the reference's reading frame, which is the default behavior. However, some consensus sequences really are out of frame relative to the reference. The indel penalties can be tuned so that RIFRAF correctly identifies true frameshifts, with a small risk of allowing some spurious ones.

### v. Multi-stage optimization

The full optimization procedure proceeds in stages, allowing RIFRAF to converge quickly by focusing on different objectives in different stages.

1. Initial stage: Do not use the reference. Propose all mutations to the consensus that appear in the pairwise alignments. Use the fixed batch, if available. If no reference was provided, stop after this step.
2. Frame correction stage: Use the full model, including the reference and reads. Propose indel candidate mutations that appearin the consensus-reference alignment. Increasingly penalize single indels in alignment of *r* and *c*. Use the fixed batch, if available.
3. Refinement stage: Propose only substitutions (no indels) to the consensus that appear in the pairwise alignments, no longer considering the reference. Use random batches, with decreasing *ρ*.

The initial stage quickly finds a good candidate consensus from the reads alone. The frame correction stage uses a reference to penalize indels that cause frame shift errors, correcting the reading frame of the template in a way that is maximally compatible with both the reads and the reference. Finally, the refinement stage ensures that the reference influences only the frame of c, and exerts no bias upon the nucleotides themselves. The final stage also fixes biases introduced by the fixed batch in the first two stages.

## III. Results

We compared `RIFRAF` to two other methods: `MAFFT` [10] followed by the standard per-column consensus, and `POA` [13] with the heaviest bundle consensus algorithm [12]. All three methods were run with and without reference-guided frame correction. `RIFRAF` natively per-forms frame correction, but only if it is given a reference sequence. To distinguish these models in this section, we refer to the model with no reference as RIFRAF_nr_, and the model with a reference as RIFRAF_ref_. `FrameBot` [24] was used for correcting results from `MAFFT` and `POA`; these are referred to as `MAFFT_FB` and `POA_FB`.

A full-length sequencing run of Pacific Biosciences SMRT sequencing on *env* from HIV-1 subtype B strain NL4-3 was used for the comparison [11]. The true sequence of NL4-3 is known, so results could be compared to the ground truth. The filtered data (available on FigShare^4^) contains 27,600 reads with expected error rate 1% or better, which were further filtered and processed as follows. To make the problem more challenging and better reveal differences between methods, very high quality sequences were excluded (expected error rate < 0.1%). Short fragments and long reads (often concatemers) were discarded by filtering out sequences 25 bases shorter or longer than the median of 2,597. PacBio reads come in random orientations, so reads were converted to their reverse complement, if necessary. Extra bases around the amplicon were removed by aligning to NL4-3 *env* without penalizing terminal gaps, then trimming terminal insertions. After preprocessing, 9,473 sequences remained, with a mean error rate of 0.0015 (the distribution of errors appears in Figure S1). All experiments were run for 1,000 trials on randomly sampled reads.

### Choice of reference

A set of reference sequences – shown in Figure S2 – were tested to investigate how frame correction accuracy deteriorates for distantly-related references. The results are shown in Figure 4. Nucleotide results from `MAFFT_FB` and `POA_FB` were both equally insensitive to the choice of reference, whereas RIFRAF_ref_′s results did degrade slightly. However, the reverse is true for the protein sequences, with RIFRAF_ref_'s performance degrading by half an amino acid on average, and the others degrading by more than one. This difference indicates that RIFRAF_ref_ not only keeps the consensus in-frame, but also makes better choices of inferring which nucleotides are truly indel errors. Finally, RIFRAF_ref_ was the most accurate, regardless of choice of reference. As expected, RIFRAF_ref_′s frame correction strategy works best with a closely related reference, but these results show that it is capable of working even with a distant reference. Except where noted, the most distant reference, B.BR, was used for the rest of the results.

**Figure 4:**
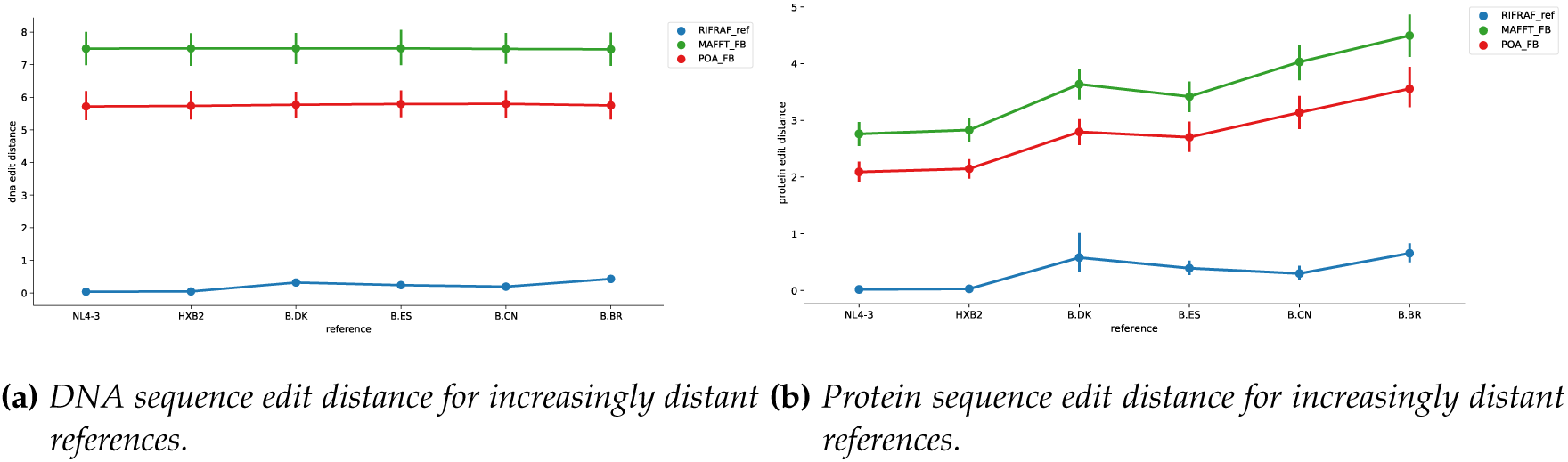
*Edit distance for increasingly distant references. All methods do better with closely-related reference, but their rate of performance degradation is important because a related reference may not always be available. Run with N* = 3 *full-length reads*.

### Number of sequences

Clusters of 2, 3, 4, 5, 6, 8, 10, 15, and 20 reads were randomly sampled for this experiment. The fraction of perfectly reconstructed consensus sequences per 1,000 trials appears in Figure 5a. For fewer than ten sequences, both versions of `RIFRAF` dominate the other corresponding methods. For instance, RIFRAF_ref_ gets over 90% correct with access to only four reads. `POA_FB` does not achieve similar results until *N* = 8, and `MAFFT_FB` does not until between *N* = 10 and 15. Interestingly, `POA`'s results actually degrade significantly for *n* > 6, but `POA_FB` continues to improve, because `POA` tends to include extra bases on the ends of the consensus sequence which are then removed by `FrameBot`. These extra bases also affect the average number of nucleotide errors (Figure 5b): for *N* = 20, `POA` averages one error per sequence, whereas all the other methods average none.

**Figure 5:**
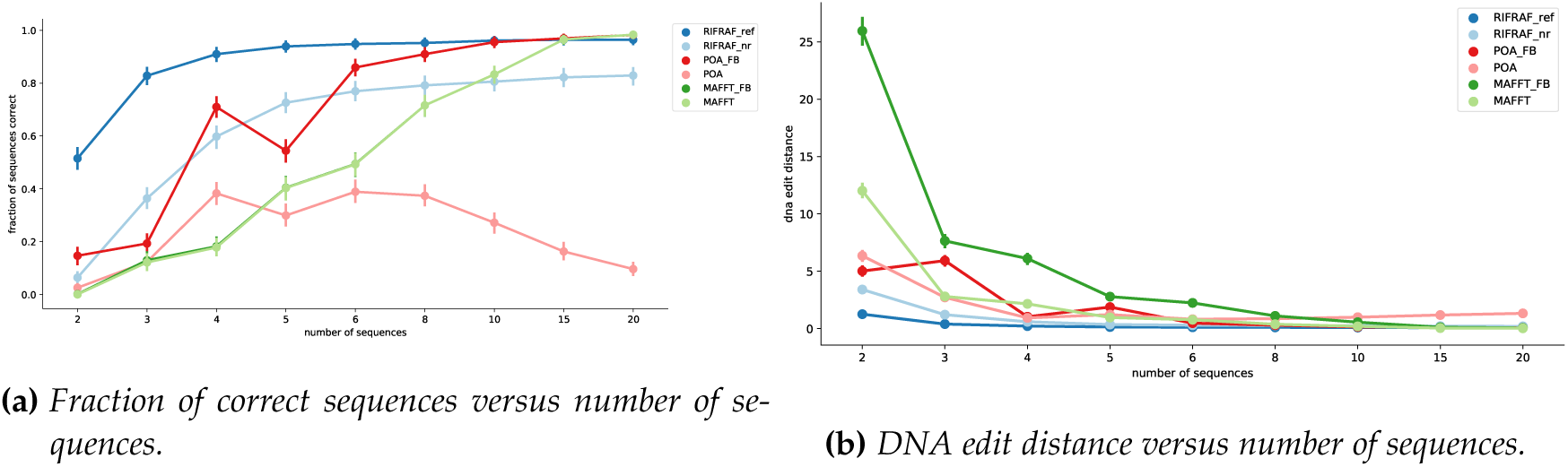
*DNA results. Fraction of correct sequences (left) and mean edit distance between the consensus and the template (right) for increasing N*.

The average number of protein errors (Figure 6a) highlights the importance of frame correction. Frame shifts cause the translated consensus sequences to differ greatly from the true protein sequence, especially for *n* < 15. For *N* = 2, fully half of each protein sequence is wrong on average, regardless of method. Even for *N* = 20, sequences from RIFRAF_nr_ and POA contain about 100 errors. On the other hand, the corrected sequences (shown in Figure 6b for clarity) contain nearly no errors for *n* > 10. RIFRAF_ref_ again performs best here, approaching zero errors even for *N* = 3.

**Figure 6:**
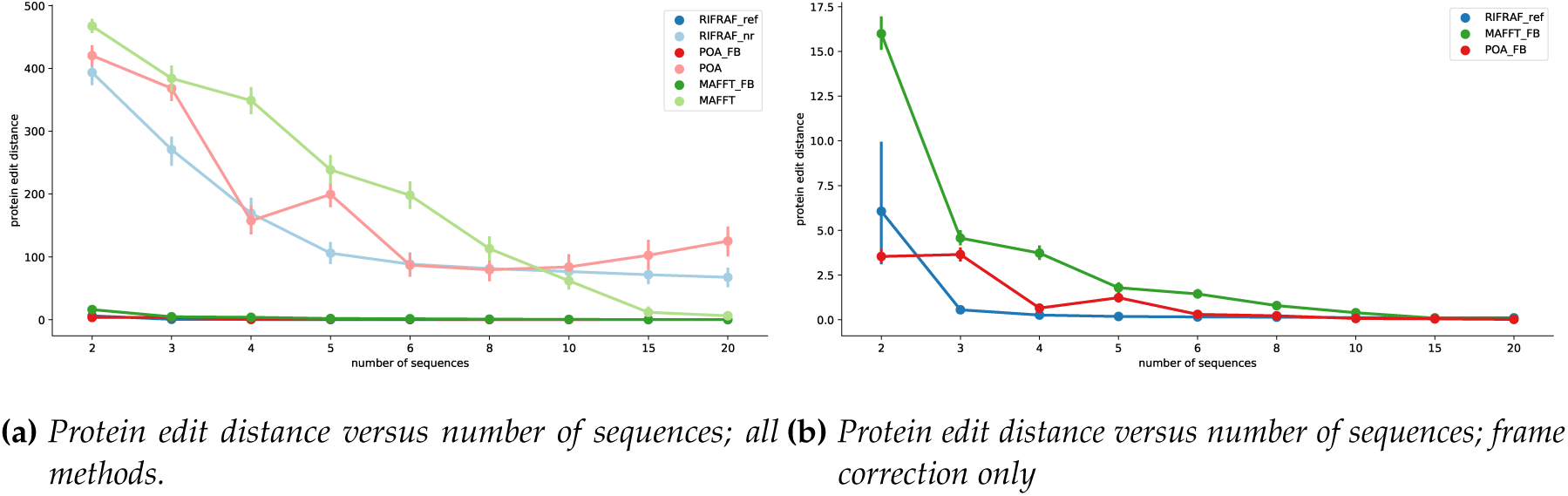
*Same results as Fig. 5, but for the translated protein sequences. The fraction of correct sequences is not reproduced, since those figures are identical. The left figure show results for all methods. The right figure show the same data, zoomed to show the frame-corrected results. Note Y axis scale*.

Interestingly, frame correction of `MAFFT` and `POA` often made the nucleotide sequences *less* accurate, whereas it improves RIFRAF_ref_. This result supports the idea that `RIFRAF`′s method of integrating frame correction into the consensus algorithm makes it more accurate by allowing all reads to inform the correction process. `FrameBot`, which only has access to a single consensus sequence, cannot use the extra information in the reads, and therefore cannot achieve the same accuracy.

Execution times appear in Figure 7. Without frame correction, all three methods are comparable for small numbers of sequences, but RIFRAF_nr_ scales better, due to its batching scheme. Frame correction adds a constant factor to all three methods’ execution times. RIFRAF_ref_′s constant factor is larger, but, because it scales better, it overtakes the others between *N* = 10 and *N* = 15.

**Figure 7:**
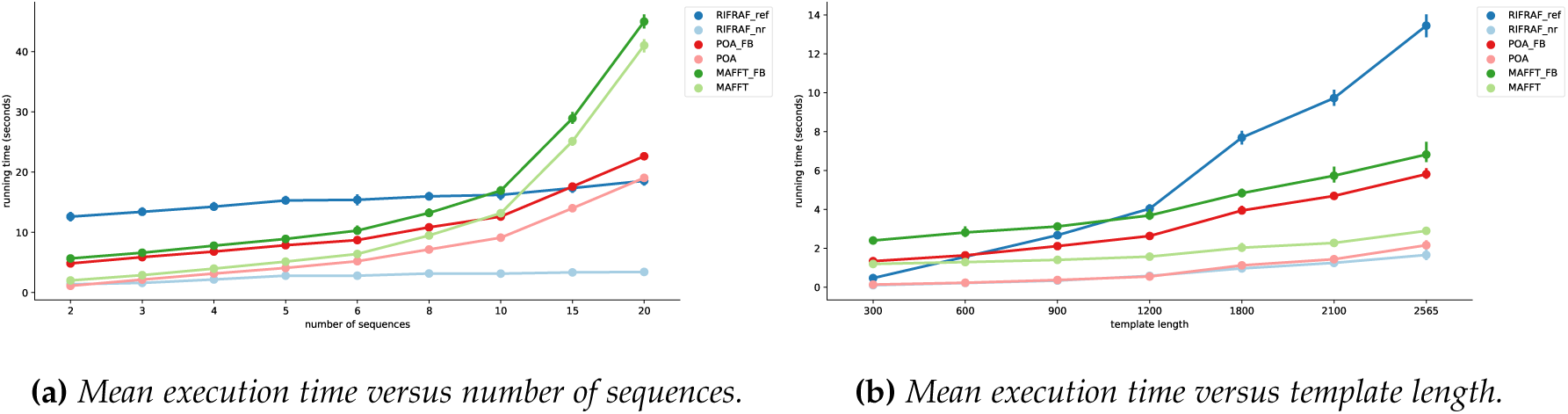
*Mean execution time, varying both number of sequences (left) and sequence length (right). Note that intervals on the x-axis are not linear*.

### Sequence length

Figure 7 also shows execution time for varying sequence lengths. For more details on this experimental setup, see SI section 2. RIFRAF_ref_ scales less well than the other methods, taking about twice as long as `MAFFT_FB` and `POA_FB` for the full-length sequences. However, it is comparable with the others at *ℓ* = 900, and faster than the others for *ℓ* < 600. This difference in speed is due to `RIFRAF`'s iterative approach, which requires recomputing parts of each pairwise alignment after every iteration.

### Detecting true frameshifts

In the other experiments, strict frameshift penalties were used to ensure the consensus stays in-frame. However, sometimes frameshifts are biologically plausible, such as in integrated (but nonfunctional) proviral Env sequences, or in the cytoplasmic tail of Env leading to a truncation, but preserving infectivity. If true frameshifts may occur in the template sequence, it may be preferable to relax this frameshift penalty. RIFRAF_ref_ can be tuned to accept frameshift indels with enough support in the reads, with only a small increase in the frequency of spurious frameshift indels. To demonstrate this, single base insertions and deletions were added to NL4-3 in both homopolymer and nonhomopolymer regions (details in SI section 3). *t_indel_* was set to 1.05, and the max frameshift indel penalty multiplier *m* varied from 0 to 12. We call an in-frame sequence a “positive”, so increasing *m* increases the false positive rate by forcing sequences with real frameshifts incorrectly into frame. To get the true positive rate, RIFRAF_ref_ was also run on the unmodified sequences. Note that while we introduce only a single true indel into our “negative” cases, the analysis is always at the whole-sequence level. We are not just detecting the presence or absence of the specific indel we introduce. Thus to achieve a high true positive and low false positive rate, RIFRAF_ref_ must successfully ignore spurious indels at *any* position in the “positive” cases, while successfully identifying the real indel we introduce in each “negative” case. The resulting ROC curves, which appear in Figure 8 for *N* ∈ 3,5,10, show that RIFRAF_ref_ can find true indels while controlling the false positive rate, using either a closely related reference or a distant one. A useful trade-off occurs for *m* = 6, which scores close to the maximum true positive rate while keeping the false positive rate close to zero.

**Figure 8:**
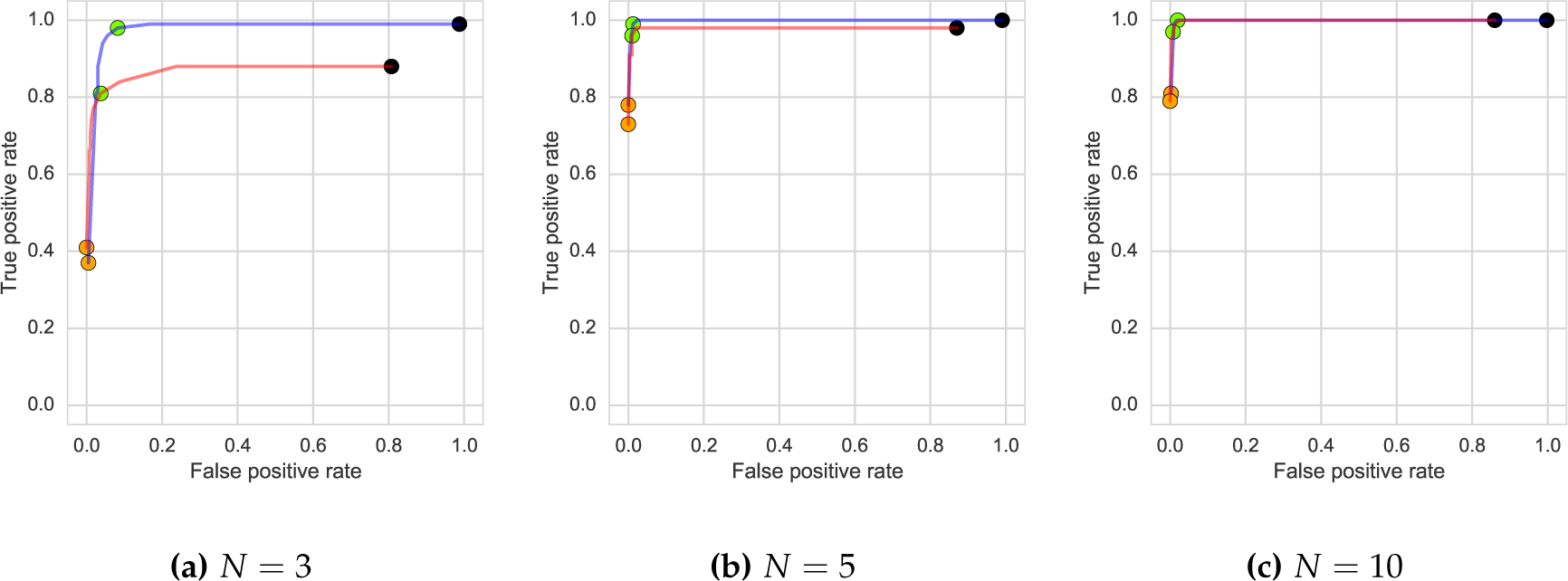
*ROC curves for true indel experiments, with max indel penalty multiplier m varying from 0 to 12. The orange point denotes results from RIFRAF_nr_, while the remainder of the curve was generated by RIFRAF_ref_. The green point corresponds to a max indel penalty multiplier m = 6. Both a related reference (HXB2, blue) and distant reference (B.BR, red) were used*.

In agreement with the accuracy results, a more closely related reference (HXB2) improved inference for *N* = 3 for this task. As expected, real homopolymer indels in homopolymer regions are harder to discriminate than non-homopolymer indels (See SI section 3 for more detailed results).

## IV. Conclusion

`RIFRAF` uses quality scores and a reference sequence to infer accurate frame-corrected consensus sequences. It can often find the correct consensus, even from small numbers of reads or with a distant reference, as shown in our experimental results. `RIFRAF` with frame correction can be slower than taking a consensus from a multiple sequence alignment, but in experiments with real SMRT sequences it finds consensus sequences that are significantly more accurate. The benefits of using a reference to reduce frameshift errors are especially apparent when comparing translated amino acid sequences, where a single frameshift causes the entire downstream sequence to be incorrect. Finally, `RIFRAF` can detect and retain true frameshifts during frame correction, and, to our knowledge, is the only method capable of this.

While `RIFRAF` performs well with distantly related reference sequences, performance is improved when using closely related references. However, when sequencing diverse populations, we note that it is always possible to first infer a set of autologous sequences from clusters or primer ID bundles that have a large number of reads, and so should be accurate. These can then be used as references to correct the reading frame of the less-represented members of the population, providing an improved accuracy over just using a more distantly related reference. We recommend using this strategy whenever possible.

`RIFRAF` can improve the ability to resolve minority variants in sequenced populations. Its ability to find results comparable to `MAFFT` with three times fewer reads will be essential for identifying minority variants in the population with greater precision. More generally, `RIFRAF` will be useful whenever an accurate consensus sequence must be inferred from a small number of full-length sequences, especially when quality scores and a reference sequence are available.

When sequencing any population, it is often advisable to sequence a clonal representative of that population first (NL4-3 *env* here), to investigate the sequencing performance for that case. We recommend using such sequence datasets to investigate the behavior of `RIFRAF` on new genes, especially if the user seeks to detect real frameshifts. To this end, we provide a Jupyter notebook that allows one to replicate the accuracy and ROC analyses from this manuscript on any clonal amplicon dataset.

`RIFRAF` will continue to be developed along multiple lines. First, the current approach for performing frame correction needs to be faster, to keep pace with the increasing volume of available sequence data. Further work needs to be done to speed it up via optimization or algorithmic advances. Possible approaches include: re-using partial alignments, speeding up alignments with k-mer seeding, and only correcting the frame of obviously problematic regions. Another improvement would include amino acid matching penalties in the reference-to-template alignment, which would allow even more distantly related reference sequences to be used, where the nucleotide homology has been completely obliterated. Another useful feature would be to infer calibrated quality scores for the consensus sequence, in order to communicate uncertain regions to the user. Finally, `RIFRAF` is extensible to other systems and sequencing technologies. In particular, we plan to investigate its behavior and tune its error model for Oxford Nanopore data, and to extend the method to support amplicons containing both non-coding and coding regions, which may contain different (potentially overlapping) reading frames.

The `RIFRAF` source code is available at https://github.com/MurrellGroup/Rifraf.jl.

## Acknowledgements

We are grateful to Sasha Murrell for copy editing.

## Footnotes

1 https://sourceforge.net/projects/poamsa/

2 https://github.com/PacificBiosciences/pbdagcon

3 https://github.com/PacificBiosciences/GenomicConsensus

4 https://doi.org/10.6084/m9.figshare.5643247

